# Bioimpedance spectroscopy assessment of skeletal muscle tissue properties after muscle damage

**DOI:** 10.64898/2026.05.21.726237

**Authors:** Jérémie Bouvier, Silvère De Freitas, Alain Letourneur, Etienne Gouraud, Alexandre Fouré

## Abstract

High intensity and unaccustomed physical activity can induce skeletal muscle damage, while even routine movements can cause similar alterations in patients with neuromuscular disease. However, reliable assessment of muscle damage using indirect markers such as muscle function evaluation, invasive measurements, or imaging techniques is difficult to implement in routine clinical follow-up and sport field settings. Bioimpedance spectroscopy appears as a promising non-invasive, easy-to-use and transportable tool to assess indirect markers of muscle damage. The aim of this study was to determine whether bioimpedance spectroscopy data are sensitive to eccentric exercise-induced muscle damage and if these potential changes mirror responses in muscle function and tissue mechanical properties. Changes in knee extensors maximal isometric contraction torque, muscle soreness, resting rigidity of the *quadriceps femoris* muscle tissue, and bioimpedance parameters at rest and during maximal isometric contraction were assessed in nine healthy males before, immediately after and in the three days following 120 maximal isokinetic eccentric contractions. Maximal contraction torque was significantly reduced during the three days following the eccentric exercise (up to -24.2%) while muscle soreness and rigidity of the *quadriceps femoris* were elevated until the second day (+494.4% and +7.6%). Changes in bioimpedance spectroscopy parameters were transiently observed at rest immediately after the damaging exercise, but not in the days that followed. Although the changes in bioimpedance parameters correlated with that of the indirect markers of muscle damage, they had already returned to baseline while functional and mechanical impairments persisted. Therefore, bioimpedance spectroscopy measurements may represent a suitable and cost-effective means of monitoring muscle fatigue.

## Introduction

Bioimpedance analysis is widely used to estimate body composition by assessing the resistive properties of biological tissues to the propagation of an alternating electrical current, particularly in athletic and clinical populations^1,2^. The biological impedance (Z), used in the predictive equation to evaluate body compartments, is derived from the resistance (R) and reactance (X_C_) of the system under investigation^3^. R primarily reflects the amount of water through which the current passes, thereby serving as an indirect marker of tissue hydration^4^. X_C_, in contrast, depends on the capacitive resistance of cells, which is determined by the structure of membrane phospholipids and proteins, making it an indirect indicator of cell integrity^4^. Furthermore, bioimpedance measurements in biological tissue are characterized by a phase shift between voltage and current signals. The angular difference between these signals, known as the phase angle (phA), is widely considered as an index of cellular health^5^.

Considering their relationship with cellular integrity, these parameters may be influenced by muscle damage. Bioimpedance could therefore represent a useful and easy-to-use tool for monitoring skeletal muscle integrity in neuromuscular disorders such as muscular dystrophies and inflammatory myopathies^6^, or to follow recovery after strenuous physical exercise. This would be particularly relevant as commonly used indirect markers of muscle damage, such as maximal voluntary force^7^ or muscle mechanical properties^8^, are harder to implement in routine clinical follow-up and sport field settings. Local decreases in both R and X_C_ have been reported in the days following muscle damage induced by *biceps brachii* eccentric contractions^9^. Moreover, injured muscles have been shown to display lower R, X_C_, and phA compared with their healthy contralateral counterparts^10^. However, using bioimpedance parameters to monitor muscle damage would require their temporal dynamics to align with those of conventional indirect markers of muscle damage. To date, only one study has investigated the relationship between these markers and bioimpedance measurements, reporting no significant associations between maximal voluntary force and phA, R, Z, or X_C_ of the exercised arm^11^. The authors attributed this lack of correlation to the narrow range of frequencies employed (*i*.*e*., 5, 50, and 250 kHz), despite the sensitivity of bioimpedance parameters to muscle damage^11^. Bioimpedance spectroscopy (BIS), which relies on the application of multiple currents across a wide frequency spectrum (1–1000 kHz), may therefore represent a more relevant tool for monitoring the muscular consequences of muscle structural alterations in ambulatory settings^12^. This method enables the characterization of parameters related to the resistance of the extracellular (R_e_) and intracellular (R_i_) compartments, estimated from the Cole modeling^5^. Consequently, fluid shifts between these compartments can be estimated using the R_e_/R_i_ ratio. BIS also enables the assessment of membrane capacitance (C_M_), reflecting the membrane’s substrate transport capacity^13^, as well as the characteristic frequency (*fc*), which corresponds to the frequency at peak X_C5_ and provides insight into the relative density of muscle tissue^14^. In only one study, BIS assessments were performed following eccentric exercise of the *biceps brachii*, revealing a local decrease in R_e_, but not in R_i_, between one and four days post-exercise^15^. However, this study only assessed R_e_ and R_i_ and did not investigate how these changes related to other indirect markers of muscle damage.

The aim of the present study was to determine whether local BIS data are altered in response to exercise-induced muscle damage (EIMD) and if these potential changes mirror adaptations in muscle functional (*i*.*e*., maximal voluntary force) and mechanical (*i*.*e*., muscle rigidity) properties. This would provide insight into the potential of BIS as a portable tool for tracking muscle tissue alterations related to muscle damage.

## Materials and Methods

### Participants

An *a priori* power analysis was performed using G*Power (v.3.1.9.7, Kiel University, Germany). We conservatively computed partial eta-squared from the available summary statistics of a previous study^15^, for the effect of EIMD on R_e_ (the main BIS parameter affected by EIMD), yielding a value of 0.195. The power analysis indicated that a sample of seven participants would be sufficient to detect such large effects in the present study, with an α of 0.05 and a power of 0.80. Thus, nine healthy recreationally active male participants (24.9 ± 3.3 years, 1.77 ± 0.05 m, 73.3 ± 7.3 kg) were recruited. None of them were engaged in regular resistance training, and they were instructed to refrain from any unfamiliar or strenuous physical activity throughout the protocol. The use of medications that could influence the experimental outcomes was prohibited, and participants were asked to maintain their usual diet while avoiding alcohol and caffeine. All participants were fully informed about the procedures and provided written consent. The study complied with the latest revision of the Declaration of Helsinki and received approval from the local ethics committee (CER-UDL_2023-09-21-007).

### Experimental design

On the first day, baseline assessments (PRE) of anthropometric characteristics, isometric maximal voluntary contraction (iMVC) torque, muscle soreness, resting rigidity, and BIS parameters were performed (Figure 1). On the same day, participants completed an eccentric exercise protocol, and the assessment session was then repeated immediately after exercise (POST), as well as one (D1), two (D2), and three (D3) days later, always at the same time of day.

**Figure 1.**
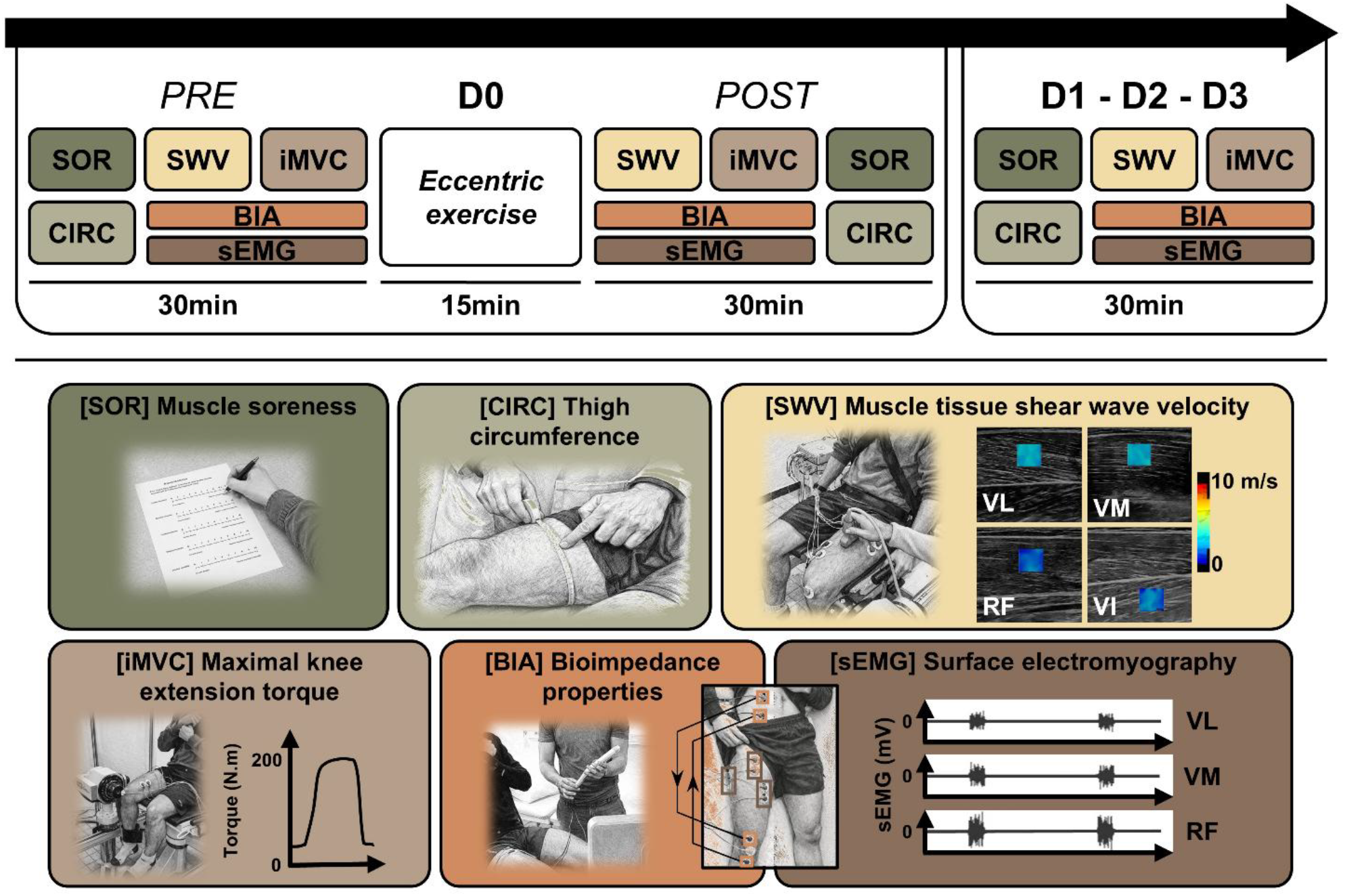
Experimental protocol and timeline of assessments. Participants were evaluated during the baseline assessment (D0), before (PRE) and immediately after (POST) an eccentric exercise, and in the three following days (from D1 to D3). At each time point, muscle soreness (SOR), thigh circumference (CIRC), resting shear wave velocity (SWV) of the vastus intermedius (VI), vastus lateralis (VL), rectus femoris (RF) and vastus medialis (VM), thigh bioimpedance parameters (BIA), and isometric maximal voluntary contraction (iMVC) torque were assessed, in this order, except for the POST session. The bioimpedance measurements were performed both at rest and during the iMVC, between two sets of emitting and receiving electrodes. Resting state was controlled by surface electromyography (sEMG). Copyrights of the photographs belong to the authors.

### Test sessions (PRE, POST, D1, D2 & D3)

#### Muscle soreness

Muscle soreness was evaluated during a passive knee extension using a visual analogue scale, ranging from ‘‘no pain’’ to ‘‘extremely painful’’.

#### Thigh circumference

Thigh circumference was measured with a tape at the midpoint of the thigh, defined as the distance between the lateral femoral condyle and the greater trochanter^16^.

#### Shear wave elastography

Muscle rigidity was assessed at rest within the *quadriceps femoris* (QF) using an Aixplorer ultrasound system (MACH30, v.2.1.0.3395, Supersonic Imagine, Aix-en-Provence, France) equipped with a linear transducer array (5–18 MHz, SuperLinear 15-8, Vermon, Tours, France). Resting muscle shear wave velocity (SWV) measurements were performed on an isokinetic dynamometer (Contrex MJ, Physiomed AG, Germany) with the hip at 70° (0° = full extension) and the knee at 100° (0° = full extension). SWV measurements were conducted on both proximal and distal sites within each head of the right QF^17^. The proximal and distal sites were defined as the 5 cm segment proximal or distal to the reference point, which was set at 50% of the thigh length for the *vastus lateralis, rectus femoris* and *vastus intermedius* and at 30% for the *vastus medialis*. The probe was carefully positioned along the muscle axis and aligned with fascicle orientation, perpendicular to the skin, while applying minimal pressure and controlling the muscle resting state through real-time visual monitoring of surface electromyography activity. Finally, the proximal and distal measurements of each QF head were averaged to produce a single representative value for the entire muscle group for each participant.

#### Bioimpedance measurements

Local (*i*.*e*., thigh) bioimpedance measurements (R, X_C_, Z and phA at 5, 50 and 200 kHz) and specific BIS parameters (C_M_, *fc*, R_e_, R_i_ and the R_e_/R_i_ ratio) were obtained using a BIS device (BiodyXpert^ZM^3, Aminogram SAS, La Ciotat, France) at rest and during iMVCs. Data were collected using two pairs of Ag/AgCl adhesive electrodes (Lessa Foam ECG, Lessa, Barcelona, Spain) placed on the shaved and alcohol-cleaned proximal and distal regions of the right QF. For the proximal region, the emitting electrode was positioned on the iliac crest and the receiving electrode at the level of the femoral head. For the distal region, the receiving and emitting electrodes were placed over the quadricipital and patellar tendons, respectively. Measurements were standardized by positioning participants on the isokinetic ergometer with the hip at 70° and the knee at 100°. For measurements during maximal contraction, participants performed an iMVC for approximately 5–6 seconds, while maintaining the same hip and knee angles.

#### Isometric maximal voluntary contractions

Following a warm-up that included twenty submaximal isometric knee extensions at a knee angle of 60°, iMVC torque was assessed at 100° using the right thigh (hip angle: 70°). At least two unilateral iMVC trials were performed, separated by a minimum rest period of 30 seconds. The highest torque value across the trials was considered for analysis.

#### Surface electromyography

The skin was shaved and cleaned with alcohol before placing pairs of Ag/AgCl surface electrodes (Foam Electrode Solid Gel 40×36 mm, Medico Electrodes, Uttar Pradesh, India) on each superficial head of the QF (*i*.*e., vastus lateralis, vastus medialis, rectus femoris*), according to the SENIAM recommendations^18^. Raw signals were recorded at 2 kHz using a PowerLab system (16/35, ADInstruments, Oxford, UK) and amplifier (FE234 Quad BioAmp, ADInstruments, Oxford, UK) and were notch- and band-pass filtered at 50 Hz and 20–400 Hz, respectively. The root mean square of surface electromyographic signals was calculated over a 150 ms moving window during resting elastography and bioimpedance measurements and normalized to the maximal value obtained during the iMVC. Recordings with a normalized root mean square above 5% were excluded from analyses to ensure passive conditions but none of the measurements exceeded this threshold.

### Eccentric exercise

Participants were seated on the isokinetic dynamometer with the hip flexed at 70° and the knee range of motion set between 20° and 100° of flexion. A brief familiarization with eccentric contractions was performed, consisting of one submaximal and two maximal efforts. Participants completed eight sets of fifteen maximal eccentric contractions at 60°/s. Between sets, participants underwent a one-minute passive recovery period. Surface electromyographic activity of QF superficial heads was also evaluated. Torque and knee angle were synchronously recorded with the PowerLab system at 2 kHz. Knee joint torque was low-pass filtered at 10 Hz and corrected for gravity. Total work was calculated for each contraction and subsequently summed for each set.

### Statistics

The reproducibility data for these measurements have already been published, with coefficients of variation ranging from 2.8 to 11.3% for bioimpedance measurements at rest, from 3.0 to 22.2% for bioimpedance measurements during iMVC, 1.4% for thigh circumference, 4.2% for iMVC torque^19^ and 2.2% for SWV^17^.

Statistical analyses were performed using R (version 4.1.1, The R Foundation for Statistical Computing). Linear mixed-effects analyses were conducted using the *lme4* R package with intercepts for participants as random effects. Visual inspection of residual plots revealed no deviations from homoscedasticity or normality, nor any violation of the linearity assumption. Absence of collinearity and influential data points was carefully checked. Bonferroni correction for multiple comparisons was applied to the post hoc analyses and associated effect sizes were computed using Cohen’s *d*. Total work was analysed with *set* as a fixed effect. iMVC torque, score of muscle soreness, thigh circumference, SWV, bioimpedance parameters (Z, phA, R and X_C_ at 5, 50 and 200 kHz as well as Re, Ri, C_M_, *fc* and R_e_/R_i_), assessed at rest and during contraction were analysed with *time* as a fixed effect.

Repeated-measures correlation was used to assess the intra-individual association between the bioimpedance parameters identified by the linear mixed models as sensitive to EIMD and other functional outcomes affected by the exercise, using the R package *rmcorr* that accounts for the non-independence of repeated observations within subjects^20^. Data are presented as mean ± standard deviation and the significance level was set at P < 0.05.

## Results

The total work performed during the eccentric exercise, iMVC torque, muscle soreness and SWV values were already reported in a previous article^17^.

### Functional responses to the eccentric exercise

The total work performed during the eccentric exercise decreased across sets (P = 0.003, Figure 2**A**), with significantly lower values in set 7 compared with set 1 (-12.7 ± 14.5%, P = 0.020, *d* = 0.726), and in set 8 compared with sets 1 and 2 (-13.4 ± 16.3%, P = 0.013, *d* = 0.718 and -12.5 ± 15.3%, P = 0.038, *d* = 0.654, respectively).

**Figure 2.**
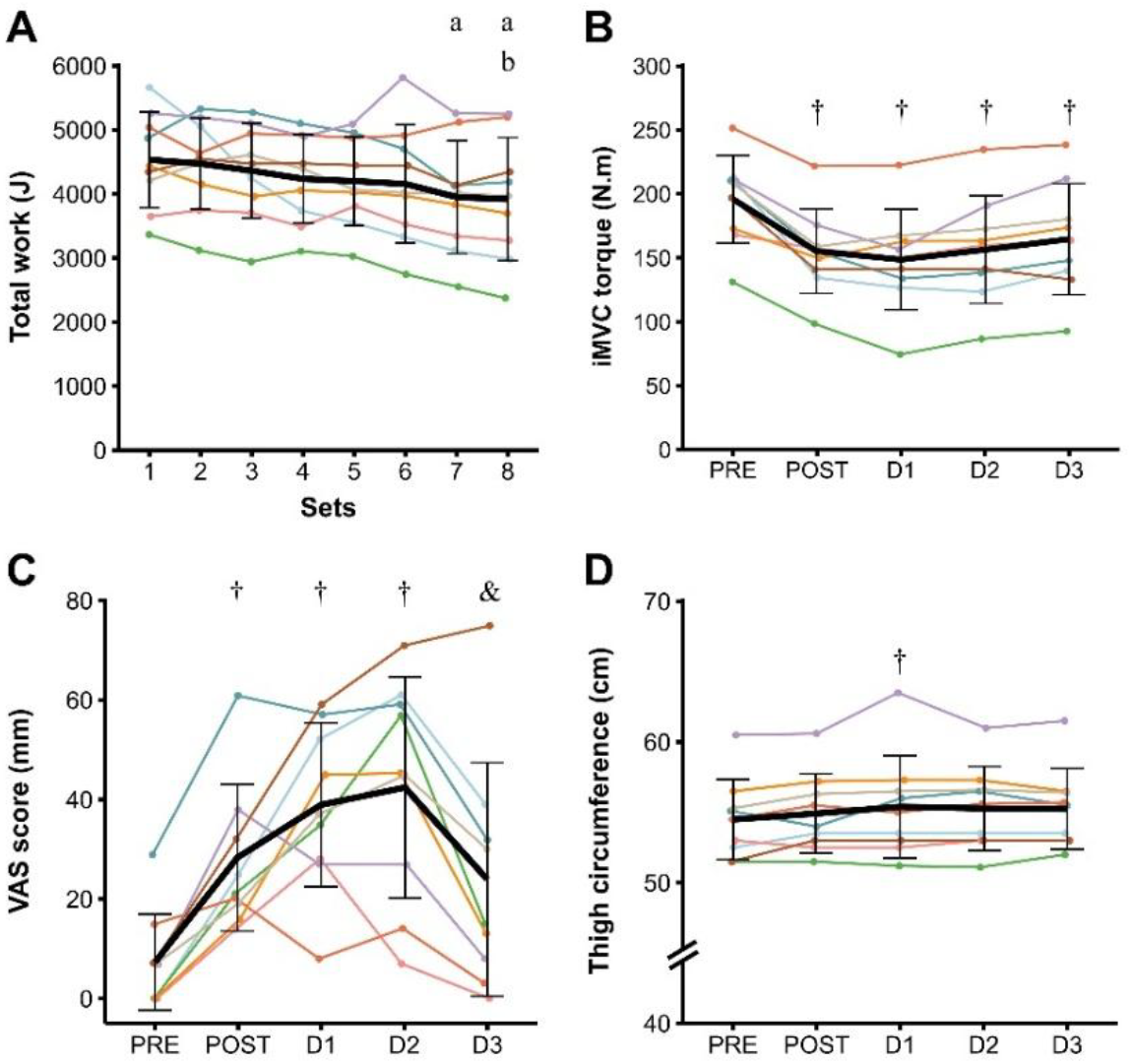
**A-**Total work performed during the eight sets of the eccentric exercise, **B-**isometric maximal voluntary contraction (iMVC) torque, **C-**muscle soreness assessed from a visual analog scale (VAS) and **D-**thigh circumference before (PRE), immediately after (POST), one (D1), two (D2) and three (D3) days after the eccentric exercise. Data are presented as mean ± standard deviation. a: significantly different from Set 1, b: significantly different from Set 2, †: significantly different from PRE, &: significantly different from D2.

iMVC torque was significantly reduced in the days following the eccentric exercise (P < 0.001, Figure 2**B**), with decreases of -20.9 ± 9.6% at POST (P < 0.001, *d* = 1.22), -24.6 ± 13.7% at D1 (P < 0.001, *d* = 1.28), -20.2 ± 14.5% at D2 (P < 0.001, *d* = 1.01), and -16.2 ± 15.1% at D3 (P < 0.001, *d* = 0.80).

Muscle soreness was significantly affected by the eccentric exercise (P < 0.001, Figure 2**C**), increasing at POST (P = 0.025, *d* = 1.61), D1 (P < 0.001, *d* = 2.33) and D2 (P < 0.001, *d* = 2.08), and then decreased at D3 (P = 0.039, *d* = 0.83, compared with D2), returning to baseline (P = 0.104, compared with PRE).

The eccentric exercise significantly increased thigh circumference (P = 0.017, Figure 2**D**), with a 1.6 ± 1.8% rise from PRE to D1 (P = 0.027, d = 0.27).

### Shear wave elastography

The SWV of the whole QF was altered by the eccentric exercise (P < 0.001, Figure 3) with significant increases observed at POST (+18.5 ± 11.9%, P < 0.001, *d* = 0.97), D1 (+8.7 ± 9.3%, P = 0.001, *d* = 0.56) and D2 (+7.6 ± 8.4%, P = 0.005, *d* = 0.48). The SWV at POST was significantly higher than at D1 (-7.7 ± 9.8%, P < 0.001, *d* = 0.53), D2 (-9.0 ± 4.1%, P < 0.001, *d* = 0.55) and D3 (-12.0 ± 5.8%, P < 0.001, *d* = 0.77).

**Figure 3.**
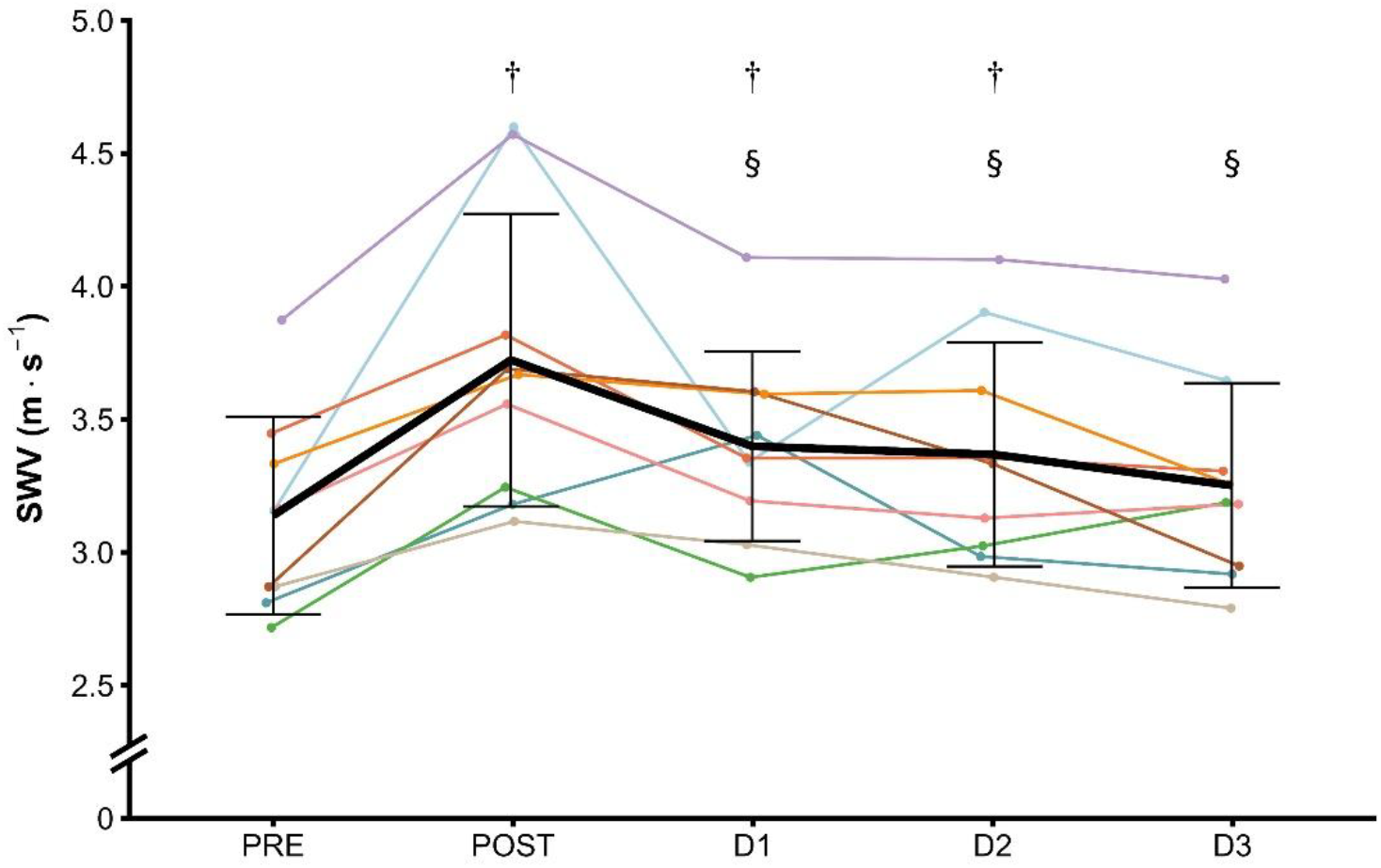
Shear wave velocity (SWV) of the quadriceps femoris before (PRE), immediately after (POST), one (D1), two (D2) and three (D3) days after the eccentric exercise. Data are presented as mean ± standard deviation. †: significantly different from PRE, §: significantly different from POST. The shear wave velocity was measured in two regions of the four quadriceps femoris heads and then averaged to obtain a representative value for the entire muscle group.

### Bioimpedance assessment

#### Thigh bioimpedance at rest

Z and R were not significantly affected by the eccentric exercise, regardless of the measurement frequency, while phA and X_C_ were altered at 5 kHz only (Table 1). phA at 5 kHz was decreased at POST compared with PRE (-16.3 ± 9.2%, P < 0.001, *d* = 1.10), D1 (-18.0 ± 11.4%, P < 0.001, d = 1.28), D2 (-14.6 ± 12.0%, P = 0.001, *d* = 0.95) and D3 (-11.3 ± 12.9%, P = 0.035, *d* = 0.81, Figure 4**A**). Similarly, X_C_ at 5 kHz was reduced at POST compared with PRE (-20.2 ± 8.7%, P < 0.001, *d* = 0.92), D1 (-21.1 ± 12.6%, P < 0.001, d = 1.06), D2 (-17.3 ± 12.2%, P = 0.002, *d* = 0.81) and D3 (-14.5 ± 16.6%, P = 0.027, *d* = 0.72, Figure 4**B**).

**Table 1.** Thigh bioimpedance measured at rest before (PRE), immediately after (POST), one (D1), two (D2), and three (D3) days after the eccentric exercise.

|  | Frequency | PRE | POST | D1 | D2 | D3 | P |
| --- | --- | --- | --- | --- | --- | --- | --- |
| Z ($\Omega$) | 5 | 66.9 ± 9.9<br>[60.4 – 73.4] | 64.0 ± 10.7<br>[57.0 – 71.0] | 66.2 ± 7.6<br>[61.2 – 71.2] | 65.7 ± 7.8<br>[60.6 – 70.8] | 66.7 ± 9.9<br>[60.2 – 73.2] | 0.355 |
| | 50 | 53.2 ± 7.4<br>[48.4 – 58.0] | 51.3 ± 8.1<br>[46.0 – 56.6] | 52.4 ± 6.2<br>[48.3 – 56.5] | 52.2 ± 5.9<br>[48.3 – 56.1] | 52.9 ± 8.2<br>[47.5 – 58.3] | 0.594 |
| | 200 | 47.0 ± 6.7<br>[42.6 – 51.4] | 44.8 ± 7.0<br>[40.2 – 49.4] | 45.9 ± 5.6<br>[42.2 – 49.6] | 45.8 ± 5.2<br>[42.4 – 49.2] | 46.4 ± 7.3<br>[41.6 – 51.2] | 0.268 |
| phA ($^\circ$) | 5 | 6.3 ± 0.9<br>[5.7 – 6.9] | 5.3 ± 1.0<sup>†</sup><br>[4.6 – 6.0] | 6.4 ± 0.8<sup>§</sup><br>[5.9 – 6.9] | 6.2 ± 1.0<sup>§</sup><br>[5.5 – 6.9] | 5.9 ± 0.6<sup>§</sup><br>[5.5 – 6.3] | **<0.001** |
| | 50 | 8.9 ± 1.1<br>[8.2 – 9.6] | 8.6 ± 1.1<br>[7.9 – 9.3] | 9.0 ± 0.5<br>[8.7 – 9.3] | 9.0 ± 0.8<br>[8.5 – 9.5] | 9.0 ± 0.6<br>[8.6 – 9.4] | 0.102 |
| | 200 | 5.0 ± 0.9<br>[4.4 – 5.6] | 4.9 ± 1.0<br>[4.2 – 5.6] | 5.0 ± 0.6<br>[4.6 – 5.4] | 5.0 ± 0.4<br>[4.7 – 5.3] | 5.1 ± 0.5<br>[4.8 – 5.4] | 0.932 |
| R ($\Omega$) | 5 | 66.5 ± 9.8<br>[60.1 – 72.9] | 63.7 ± 10.6<br>[56.8 – 70.6] | 65.8 ± 7.6<br>[60.8 – 70.8] | 65.3 ± 7.7<br>[60.3 – 70.3] | 66.4 ± 9.8<br>[60.0 – 72.8] | 0.389 |
| | 50 | 52.5 ± 7.2<br>[47.8 – 57.2] | 50.8 ± 8.0<br>[45.6 – 56.0] | 51.8 ± 6.1<br>[47.8 – 55.8] | 51.6 ± 5.8<br>[47.8 – 55.4] | 52.2 ± 8.2<br>[46.8 – 57.6] | 0.608 |
| | 200 | 46.8 ± 6.7<br>[42.4 – 51.2] | 44.7 ± 7.0<br>[40.1 – 49.3] | 45.7 ± 5.6<br>[42.0 – 49.4] | 45.7 ± 5.2<br>[42.3 – 49.1] | 46.2 ± 7.3<br>[41.4 – 51.0] | 0.267 |
| X<sub>C</sub> ($\Omega$) | 5 | 7.4 ± 1.5<br>[6.4 – 8.4] | 5.9 ± 1.6<sup>†</sup><br>[4.9 – 6.9] | 7.4 ± 1.2<sup>§</sup><br>[6.6 – 8.2] | 7.1 ± 1.4<sup>§</sup><br>[6.2 – 8.0] | 6.9 ± 0.9<sup>§</sup><br>[6.3 – 7.5] | **<0.001** |
| | 50 | 8.3 ± 1.7<br>[7.2 – 9.4] | 7.7 ± 1.7<br>[6.6 – 8.8] | 8.2 ± 1.1<br>[7.5 – 8.9] | 8.1 ± 1.2<br>[7.4 – 9.0] | 8.3 ± 1.3<br>[7.5 – 9.1] | 0.273 |
| | 200 | 4.1 ± 1.1<br>[3.4 – 4.8] | 3.8 ± 1.1<br>[3.1 – 4.5] | 4.0 ± 0.7<br>[3.5 – 4.5] | 4.0 ± 0.6<br>[3.6 – 4.4] | 4.1 ± 0.8<br>[3.6 – 4.6] | 0.732 |
| BIS parameters | R<sub>e</sub> ($\Omega$) | 72.1 ± 11.1<br>[64.8 – 79.4] | 67.4 ± 11.6<br>[59.8 – 75.0] | 71.0 ± 8.3<br>[65.6 – 76.4] | 70.3 ± 8.7<br>[64.6 – 76.0] | 70.8 ± 10.2<br>[64.1 – 77.5] | 0.119 |
| | R<sub>i</sub> ($\Omega$) | 125.8 ± 21.8<br>[111.6 – 140.0] | 126.4 ± 24.7<br>[110.3 – 142.5] | 122.5 ± 19.9<br>[109.5 – 135.5] | 124.6 ± 18.4<br>[112.6 – 136.6] | 126.8 ± 26.7<br>[109.4 – 144.2] | 0.716 |
| | C<sub>M</sub> (nF) | 42.4 ± 8.0<br>[37.2 – 47.6] | 36.1 ± 9.1<sup>†</sup><br>[30.2 – 42] | 40.1 ± 7.6<br>[35.1 – 45.1] | 41.1 ± 10.5<br>[34.2 – 48.0] | 38.1 ± 8.2<br>[32.7 – 43.5] | **0.031** |
| | fc (kHz) | 19.7 ± 2.3<br>[18.2 – 21.2] | 23.8 ± 3.4<sup>†</sup><br>[21.6 – 26.0] | 20.7 ± 2.5<sup>§</sup><br>[19.1 – 22.3] | 20.8 ± 3.4<sup>§</sup><br>[18.6 – 23.0] | 22.0 ± 2.2<br>[20.6 – 23.4] | **<0.001** |
| | R<sub>e</sub>/R<sub>i</sub> ($\Omega$) | 0.58 ± 0.10<br>[0.51 – 0.65] | 0.54 ± 0.11<br>[0.47 – 0.61] | 0.59 ± 0.08<br>[0.54 – 0.64] | 0.57 ± 0.09<br>[0.51 – 0.63] | 0.57 ± 0.06<br>[0.53 – 0.61] | 0.331 |

Values are presented as mean ± standard deviation [confidence interval 95%]. Each *p*-value corresponds to the main effect of time on the associated variable. Significant differences are indicated by bold *p*-values. †: significantly different from PRE, §: significantly different from POST. Z: bioimpedance, phA: phase angle, R: resistance, X<sub>C</sub>: reactance, BIS: bioimpedance spectroscopy, R<sub>e</sub>: extracellular resistance, R<sub>i</sub>: intracellular resistance, C<sub>M</sub>: membrane capacitance and fc: characteristic frequency.

**Figure 4.**
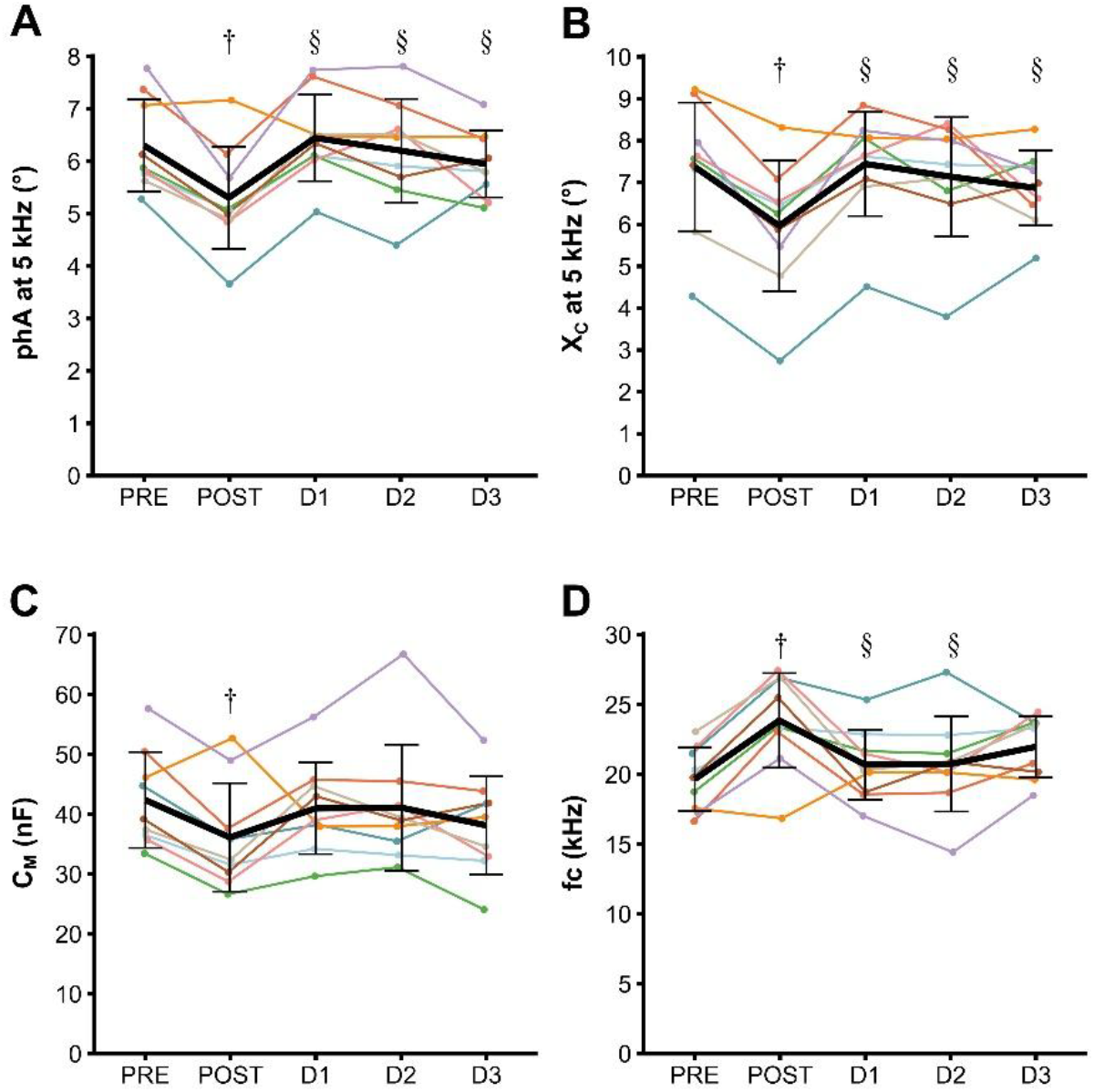
**A-**Phase angle (phA) at 5 kHz, **B-**reactance (X_C_) at 5 kHz, **C-**membrane capacitance (C_M_) and **D-**characteristic frequency (fc) measured before (PRE), immediately after (POST), one (D1), two (D2) and three (D3) days after the eccentric exercise. Data are presented as mean ± standard deviation. †: significantly different from PRE, §: significantly different from POST.

Regarding the thigh BIS parameters measured at rest, only C_M_ and *fc* were modified after the eccentric exercise (Table 1). C_M_ was reduced at POST compared with PRE (-15.1 ± 11.7%, P = 0.049, *d* = 0.73, Figure 4**C**) while *fc* was higher at POST compared with PRE (+21.4 ± 11.7%, P < 0.001, *d* = 1.45), D1 (+16.1 ± 17.2%, P = 0.006, *d* = 1.06) and D2 (+16.7 ± 19.9%, P = 0.007, *d* = 0.91, Figure 4**D**).

#### Thigh bioimpedance during contraction

None of the bioimpedance parameters (*i*.*e*. Z, R, X_C_ and phA) measured during contraction were affected by the eccentric exercise (Supplementary table 1). Similarly, the BIS parameters (*i*.*e*. R_e_, R_i_, C_M_, *fc* and R_e_/R_i_) measured during the iMVC remained unchanged (Supplementary table 1).

### Correlation analyses

Repeated-measures correlations revealed significant negative associations across time points between SWV and Xc at 5 kHz (P = 0.004, r = -0.461, Figure 5F), phA at 5 kHz (P = 0.002, r = -0.503, Figure 5E) and C_M_ (P = 0.017, r = -0.391, Figure 5C), as well as between iMVC and *fc* (P = 0.028, r = -0.362, Figure 5B). SWV also displayed a positive correlation with *fc* (P = 0.002, r = 0.487, Figure 5D).

**Figure 5.**
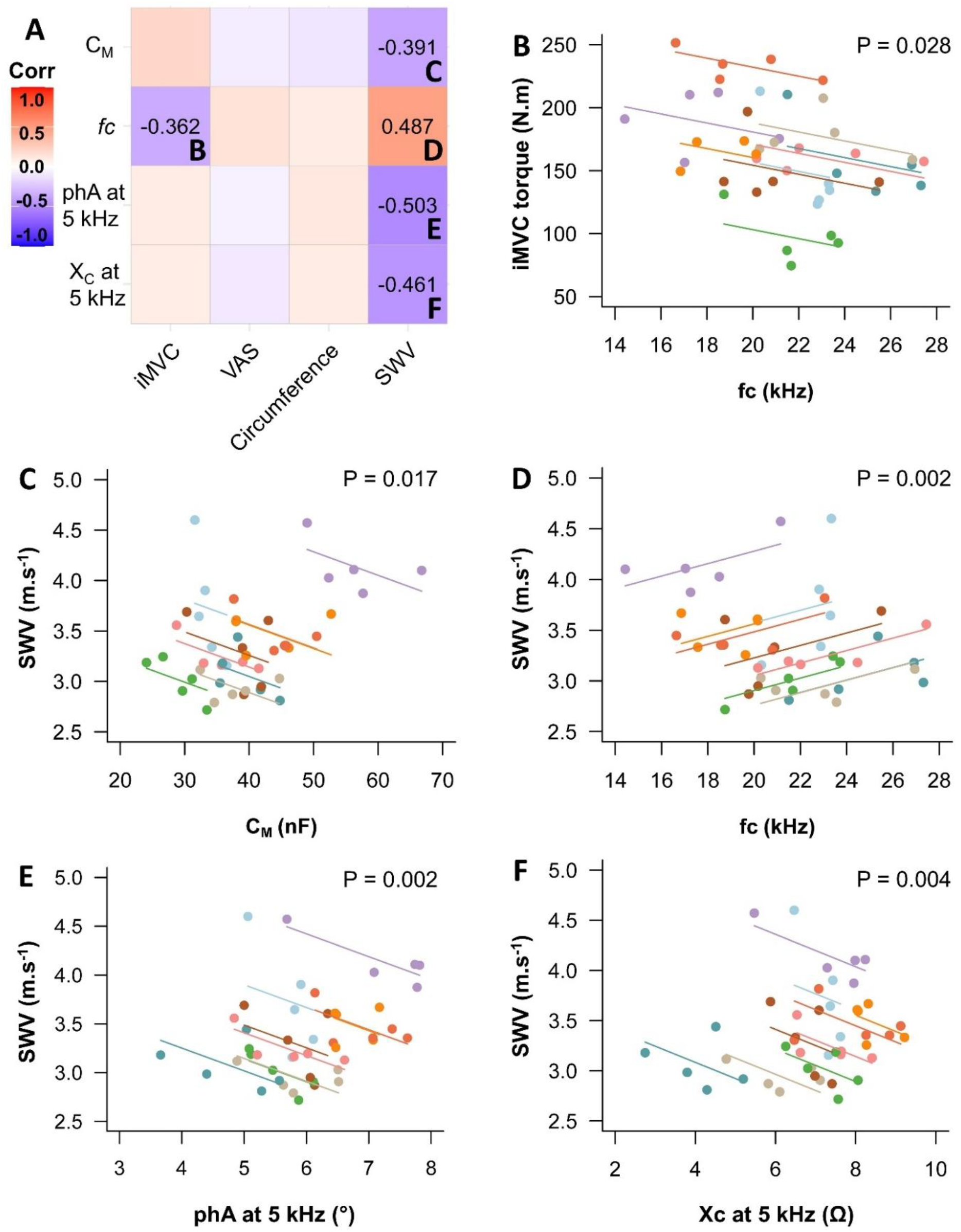
Correlations between bioimpedance measurements and indirect markers of muscle damage. Only the bioimpedance parameters identified by the linear mixed models as sensitive to eccentric exercise were compared with other functional outcomes affected by the exercise. Panel **A** depicts all the correlations tested while the correlations concerning the characteristic frequency were represented in panels **B** and **D**, membrane capacitance in panel **C**, phase angle in panel **E** and reactance in panel **F**. In panel **A**, correlation coefficients are only presented for the significant correlations. C_M_: membrane capacitance, fc: characteristic frequency, phA: phase angle, X_C_: reactance, iMVC: isometric maximal voluntary contraction, VAS: visual analogue scale and SWV: shear wave velocity.

## Discussion

The aim of the present study was to determine whether BIS data are altered following eccentric EIMD and whether any such changes reflect adaptations in functional and mechanical properties. Our results show that classical bioimpedance and spectroscopic outcomes are altered directly after eccentric exercise. In addition, BIS parameters correlated with indirect markers of EIMD during the period following eccentric exercise.

Indirect markers of muscle damage including soreness, iMVC and SWV followed the typical kinetics reported in previous EIMD studies^7,8,21,22^. A decrease in phA and X_C_ (5 kHz only) was observed at POST compared with all other time points. Because current is less likely to travel through the cell membrane at this frequency, these decreases may mainly reflect modifications in the extracellular compartments^23^. The lower phA and X_C_ at POST also suggest a decrease in sarcolemmal excitability^24^ and/or sarcolemma disruptions induced by the eccentric exercise^25^. A similar reduction was reported immediately after eccentric elbow flexors exercise^11^, but in that study the decrease persisted up to 7 days, whereas we observed a return to baseline by D1. Other studies also reported inconsistent findings, with decreases in phA and X_C_ appearing only from D1^10^ or D3^9^ while it was unaffected over this time frame in our study. Such discrepancies likely reflect methodological differences, including measurement frequencies (10–100 kHz in Freeborn et al.^6^, 50 kHz in Nescolarde et al.^10^, 5–250 kHz in Yamaguchi et al.^10^) and measurement sites (localized on the arm in Freeborn et al.^6^ and in Yamaguchi et al.^10^ and on the thigh in Nescolarde et al.^10^), preventing a clear characterization of eccentric EIMD from bioimpedance assessments.

To address such variability in measurement frequency, BIS enables concurrent assessment across multiple frequencies. R_i_ was unchanged in our study, as for the study from Shiose et al.^15^, the only previous BIS study performed after eccentric exercise. However, Shiose et al. reported a 27% decrease in R_e_ that was not observed in the present study. This discrepancy may stem from the muscles investigated, as upper-limb muscles are generally more susceptible to EIMD than lower-limb muscles and BIS measurements may differ between limbs^26^. Moreover, C_M_ decreased and *fc* increased at POST in our study, both being BIS outcomes not evaluated by Shiose et al.^15^. Decreases in C_M_ and *fc* may reflect a reduction in cell membrane integrity and/or capacity to store electrical charges^3^. However, the cell-membrane integrity disruptions caused by EIMD are particularly apparent two to four days after the exercise^27^. Therefore, it seems unlikely that the modification in C_M_ and *fc* at POST reflect alterations in cell membrane integrity. These changes might rather reflect acute fatigue after the eccentric contractions, given that their time course is very similar to the modulation of neuromuscular parameters related to fatigue and measured from twitch interpolation during the acute phase of EIMD^28^. Notably, exercise is known to induce transient changes in sarcolemmal excitability^24^ and/or structural disruption^25^, which could explain the decrease in C_M_ and increase in *fc* at POST, suggesting BIS may provide fatigue-related information^29^. Any mechanistic interpretation remains however speculative, as no previous study assessed BIS outcomes in conjunction with fatigue parameters immediately after exercise.

Contrary to our hypothesis, C_M_ and *fc* returned to baseline by D1. Based on their link to cellular integrity, we expected more prolonged changes, similar to the time course of EIMD recovery. Ultimately, *fc* was correlated with changes in iMVC, the most reliable indirect marker of EIMD^7^. Additionally, the four bioimpedance parameters that showed sensitivity to the eccentric exercise were correlated with SWV, exhibiting changes similar to that of muscle mechanical properties in response to EIMD. However, the correlation between *fc* and iMVC was weak (r = -0.362), and all bioimpedance parameters were associated with SWV as they, like SWV, changed markedly at POST (without showing sensitivity to EIMD from D1 to D3). Taken together with complementary linear regression analyses (Supplementary Figure 1), our results indicate that BIS sensitivity may not be sufficient to accurately evaluate EIMD, when measurement is performed at the whole muscle level.

On the opposite to the rest condition, bioelectrical impedance measurements during contraction were unaffected by EIMD. This may reflect their slightly lower reproducibility compared with resting measurements^19^ and/or a reduced sensitivity to EIMD confined to the thigh.

Although the sample size was relatively small, the *a priori* power analysis indicated sufficient power to detect the large magnitude of changes in bioimpedance parameters expected in response to EIMD. However, smaller effects may have remained undetected. Another limitation of this study is the absence of a control condition. Nevertheless, the high reproducibility of our measurements^19^ provides confidence that such changes could not have occurred in the absence of tissue alterations. Another limitation could be the electrode placement used in our study which may lower the sensitivity of BIS parameters to EIMD by evaluating the whole thigh, unlike other studies which focused on the injury location^10^ or smaller muscles, like the elbow flexors^9,11^.

### Practical applications

Our results indicate that BIS parameters (C_M_ and *fc*) are altered immediately following eccentric exercise. However, they return to baseline by the next day, while changes in indirect markers of EIMD persist, highlighting the moderate sensitivity of BIS to detect muscle damage when using the selected electrode configuration to assess the whole thigh. Consequently, further study should focus on the standardization of electrodes placement to improve the use of electrical impedance myography in field settings or clinical routine follow-up. Nevertheless, the observed alterations in BIS parameters immediately after eccentric contractions underscore the potential of this tool as a cost-effective means to quantify muscle fatigue, provided that a direct relationship between post-exercise BIS changes and the determinants of fatigue can be established. Future studies using graded fatigue protocols may help clarify the relationship between acute post-exercise force loss and changes in bioimpedance and BIS parameters, which may contribute to the identification of bioimpedance-based biomarkers of fatigue, especially in patients with neuromuscular disease.

## Conclusions

Although some variations in BIS parameters were weakly correlated to changes in muscle mechanical properties assessed by shear wave elastography, the present study, based on the parallel assessment of these parameters alongside classical markers of EIMD, indicates that BIS could be a suitable tool for field-based and routine clinical evaluations of muscle fatigue.

## Abbreviations

BIS: Bioimpedance spectroscopy
C_M_: Membrane capacitance
D0, D1, D2, D3: day of the eccentric exercise, one, two and three days after
EIMD: Exercise-induced muscle damage
*fc*: Characteristic frequency
iMVC: Isometric maximal voluntary contraction
phA: Phase angle
POST: Immediately after eccentric exercise
PRE: before eccentric exercise
QF: *Quadriceps femoris*
R: Resistance
R_e_: Extracellular resistance
R_i_: Intracellular resistance
R_inf_: resistance of the combined extracellular and intracellular compartments
SWE: Shear wave elastography
SWV: Shear wave velocity
X_C_: Reactance
Z: Bioimpedance

## Acknowledgments

The authors would like to express their gratitude to all volunteers for their participation in this study.

## Data availability statement

The data that supports the findings of this study are available from the corresponding author upon reasonable request.

## Ethics approval statement

The study complied with the latest revision of the Declaration of Helsinki and received approval from the local ethics committee (CER-UDL_2023-09-21-007).

## Patient consent statement

All participants were fully informed about the procedures and provided written consent.

