## Supplemental files for "Bioimpedance spectroscopy assessment of skeletal muscle tissue properties after muscle damage"

**Supplementary table 1** *Thigh bioimpedance measured during contraction before (PRE), immediately after (POST), one (D1), two (D2), and three (D3) days after the eccentric exercise.*

| | Frequency | PRE | POST | D1 | D2 | D3 | P |
|---|---|---|---|---|---|---|---|
| Z (Ω) | 5 | 66.4 ± 10.1 [59.8 – 73.0] | 63.7 ± 10.5 [56.8 - 70.6] | 65.8 ± 10.1 [59.2 - 72.4] | 64.2 ± 9.4 [58.1 - 70.3] | 64.5 ± 11.0 [57.3 - 71.7] | 0.672 |
| | 50 | 52.6 ± 7.5 [47.7 - 57.5] | 50.3 ± 8.2 [44.9 - 55.7] | 51.3 ± 7.8 [46.2 - 56.4] | 50.2 ± 6.7 [45.8 - 54.6] | 50.7 ± 8.9 [44.9 - 56.5] | 0.575 |
| | 200 | 45.8 ± 6.3 [41.7 - 49.9] | 44.7 ± 6.9 [40.2 - 49.2] | 44.9 ± 6.0 [41.0 - 48.8] | 43.7 ± 6.0 [39.8 - 47.6] | 44.7 ± 8.1 [39.4 - 50] | 0.607 |
| phA (°) | 5 | 6.5 ± 1.1 [5.8 - 7.2] | 5.8 ± 1.7 [4.7 - 6.9] | 6.5 ± 1.0 [5.8 - 7.2] | 6.4 ± 0.7 [5.9 - 6.9] | 6.2 ± 1.0 [5.5 - 6.9] | 0.413 |
| | 50 | 8.7 ± 1.6 [7.7 - 9.7] | 8.4 ± 1.1 [7.7 - 9.1] | 9.8 ± 1.4 [8.9 - 10.7] | 9.1 ± 1.6 [8.1 - 10.1] | 9.1 ± 1.6 [8.1 - 10.1] | 0.064 |
| | 200 | 4.4 ± 1.6 [3.4 - 5.4] | 4.8 ± 0.8 [4.3 - 5.3] | 5.2 ± 1.3 [4.4 – 6.0] | 4.2 ± 1.7 [3.1 - 5.3] | 4.7 ± 1.6 [3.7 - 5.7] | 0.460 |
| R (Ω) | 5 | 65.9 ± 10.0 [59.4 - 72.4] | 63.4 ± 10.5 [56.5 - 70.3] | 65.3 ± 10.0 [58.8 - 71.8] | 63.8 ± 9.3 [57.7 - 69.9] | 64.1 ± 11.0 [56.9 - 71.3] | 0.704 |
| | 50 | 52.0 ± 7.3 [47.2 - 56.8] | 49.7 ± 8.1 [44.4 – 55.0] | 50.5 ± 7.7 [45.5 - 55.5] | 49.5 ± 6.5 [45.3 - 53.7] | 50.1 ± 8.7 [44.4 - 55.8] | 0.554 |
| | 200 | 45.7 ± 6.3 [41.6 - 49.8] | 44.6 ± 6.9 [40.1 - 49.1] | 44.7 ± 5.9 [40.8 - 48.6] | 43.5 ± 6.0 [39.6 - 47.4] | 44.5 ± 8.0 [39.3 - 49.7] | 0.597 |
| $X_C$ (Ω) | 5 | 7.6 ± 1.8 [6.4 - 8.8] | 6.4 ± 2.0 [5.1 - 7.7] | 7.5 ± 1.6 [6.5 - 8.5] | 7.1 ± 1.1 [6.4 - 7.8] | 6.9 ± 1.1 [6.2 - 7.6] | 0.067 |
| | 50 | 8.0 ± 2.1 [6.6 - 9.4] | 7.3 ± 1.5 [6.3 - 8.3] | 8.7 ± 2.0 [7.4 – 10.0] | 8.0 ± 2.2 [6.6 - 9.4] | 8.1 ± 2.2 [6.7 - 9.5] | 0.243 |
| | 200 | 3.5 ± 1.4 [2.6 - 4.4] | 3.8 ± 0.9 [3.2 - 4.4] | 4.1 ± 1.4 [3.2 – 5.0] | 3.2 ± 1.5 [2.2 - 4.2] | 3.8 ± 1.6 [2.8 - 4.8] | 0.471 |
| BIS parameters | $R_e$ (Ω) | 70.9 ± 11.1 [63.6 - 78.2] | 68.2 ± 11.2 [60.9 - 75.5] | 70.0 ± 11.3 [62.6 - 77.4] | 68.1 ± 10.5 [61.2 - 75] | 68.4 ± 12.1 [60.5 - 76.3] | 0.639 |
| | $R_i$ (Ω) | 126.0 ± 22.0 [111.6 - 140.4] | 120.8 ± 24.1 [105.1 - 136.5] | 117.0 ± 10.5 [110.1 - 123.9] | 120.8 ± 13.9 [111.7 - 129.9] | 122.4 ± 23.0 [107.4 - 137.4] | 0.363 |
| | $C_M$ (nF) | 46.9 ± 9.2 [40.9 - 52.9] | 46.4 ± 14.8 [36.7 - 56.1] | 46.3 ± 9.4 [40.2 - 52.4] | 44.0 ± 11.8 [36.3 - 51.7] | 49.1 ± 16.3 [38.5 - 59.7] | 0.908 |
| | *fc* (kHz) | 17.9 ± 2.9 [16.0 - 19.8] | 19.6 ± 4.0 [17.0 - 22.2] | 19.1 ± 3.5 [16.8 - 21.4] | 20.3 ± 4.3 [17.5 - 23.1] | 18.5 ± 4.1 [15.8 - 21.2] | 0.675 |
| | $R_e/R_i$ (Ω) | 0.57 ± 0.11 [0.50 - 0.64] | 0.58 ± 0.10 [0.51 - 0.65] | 0.60 ± 0.10 [0.53 - 0.67] | 0.56 ± 0.07 [0.10 – 1.02] | 0.56 ± 0.08 [0.04 - 1.08] | 0.450 |

*Values are presented as mean ± standard deviation [confidence interval 95%]. Each p-value corresponds to the main effect of time on the associated variable. Z: bioimpedance, phA: phase angle, R: resistance, $X_C$: reactance, BIS: bioimpedance spectroscopy, $R_e$: extracellular resistance, $R_i$: intracellular resistance, $C_M$: membrane capacitance and fc: characteristic frequency.*

**Supplementary Figure 1** *Linear regression analyses between changes in EIMD markers (isometric maximal voluntary contraction [iMVC] torque, muscle soreness assessed from a visual analogue scale [VAS], thigh circumference [CIRC] and shear wave velocity [SWV]) and changes in the four BIA parameters responsive to eccentric exercise (phA and $X_C$ at 5 kHz,* fc *and $C_M$) from the baseline assessment (PRE) to one, two and three days after exercise (D1, D2 and D3).*

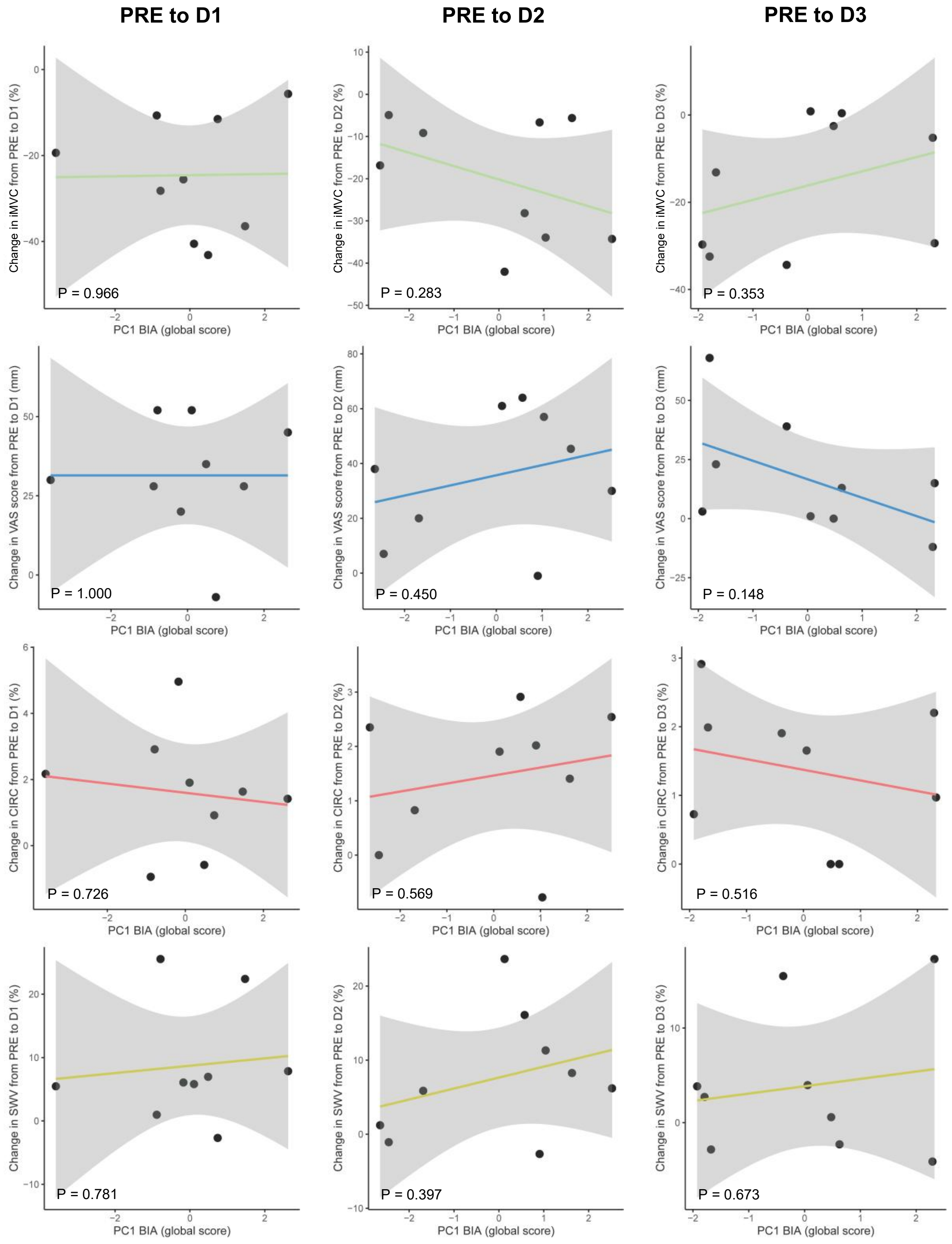

To further explore the potential relevance of BIS in relation to EIMD, linear regression analyses were performed between EIMD markers (at time points where EIMD was likely present) and the four BIA parameters responsive to eccentric exercise. These parameters were combined using a principal component analysis (PCA) to reduce collinearity among BIA variables and limit the risk of model overfitting while retaining their shared physiological information. These complementary analyses revealed no association between the BIA-derived PCs (proportion of variance explained by: PCA PRE to D1: 75.3%; PCA PRE to D2: 84.8%; PCA PRE to D3: 66.2%) and changes in EIMD markers from PRE to D1, PRE to D2, or PRE to D3 (all $P > 0.05$; see Figure above), further highlighting the low sensitivity of BIA to EIMD.
